# Honeycomb-inspired SERS nano-bowls for rapid capture and analysis of extracellular vesicles and liposomes in suspension

**DOI:** 10.1101/2023.05.18.541353

**Authors:** Sathi Das, Jean-Claude Tinguely, Sybil Akua Okyerewa Obuobi, Eduarda M. Guerreiro, Natasa Skalko-Basnet, Omri Snir, Kanchan Saxena, Balpreet Singh Ahluwalia, Dalip Singh Mehta

## Abstract

Nanoscale carriers such as liposomes and extracellular vesicles (EVs) are readily being explored for personalized medicine or disease prediction and diagnostics, respectively. Owing to their small size, such nanocarriers can undergo endocytosis or exocytosis, providing means to either transport cargo to the cells (liposomes) or to serve as a biomarker (EVs). When looking at current analysis methods, there is a growing need for detailed characterization of the content and composition of such nanocarriers in their natural state in aqueous media. This can be achieved through surface-enhanced Raman spectroscopy (SERS), which provides a molecular fingerprint of the analytes while reducing the detection limit. In this paper, we utilize a nano-structured SERS substrate to study different bio-nanoparticles such as liposomes, EVs and DNA nanogel in suspension. A silver-coated polydimethylsiloxane (PDMS) film-based honeycomb shaped nano-bowl surface passively traps and reduces the mobility of the nanosized bio-particles, improving the intensity and the reproducibility of the SERS signal. FDTD simulations are used for substrate geometry optimization, and a detection limit of 10^−15^ M is demonstrated for Rhodamine 6G (R6G). The potential of the proposed SERS nano-bowl is shown through distinct spectral features following surface-(polyethylene glycol) and bilayer-(cholesterol) modification of empty liposomes. For DNA nanogels, the characterization of highly crosslinked DNA specimens exhibits enhanced peaks for nitrogenous bases, sugar, and phosphate groups. EVs isolated from various cells provided spectral signatures of specific protein content, lipid components, and nucleic acids. Concluding, the findings of the spectral signatures of a wide range of molecular complexes and chemical morphology of bio-membranes in their natural state highlight the possibilities of using SERS as a sensitive and instantaneous characterization alternative.

## Introduction

Biological macromolecules such as lipids and nucleic acids have drawn attention as biocompatible materials in the development of novel therapeutics and diagnostic devices.^1^ The application of nanosized biological relevant particles fabricated from these biological materials (e.g., liposomes, extracellular vesicles and DNA nanogels) are rapidly expanding against non-communicable (e.g., cancers) and communicable (e.g., infections) diseases.^2-4^ Liposomes are nanosized artificial vesicles, composed of a lipid bilayer that can encapsulate drugs or other target specific molecules. Owing to their small size, parenterally administered liposomes can circulate until they reach the target cells and bind with the target directly or by endocytosis. Consequently, liposomes are finding an increased role within drug delivery and disease treatment by allowing drugs to be directed to specific cells or tissues in the body. The successful delivery of liposomes to the target cell is known to be influenced by their size. However, current studies demonstrate how surface modifications improve target specificity and the impact of bilayer modifications on guarding the stability of liposomes enroute to their target. To this end, it is essential to characterize the chemical fingerprint of liposomes to ensure that key modifications that influence their functionalities remain intact in vivo.^5,6^

Extracellular vesicles (EVs) are nanobodies confined by lipid bilayer membranes which are naturally produced by all kind of cells, and carry diverse biochemical loads of proteins, lipids, and nucleic acids (i.e., RNA and DNA).^7^ EVs display a biological complexity and diverse heterogeneity according to their origin. Adequate analysis of the composition and biochemical load of EVs may thus provide critical information towards their exploitation for personalized medicine, disease prediction and diagnosis.^8,9^

The high programmability of DNA has enabled the self-assembly of a variety of DNA nanostructures (e.g., DNA nanogels) that have controllable size, morphology and which can modulate their surface chemistry via precise display of ligands to target specific cells and tissues.^10^ Since different approaches are used in designing these nanocarriers (e.g., thermal hybridization, enzyme initiated or chemical functionalization), confirming the accurate assembly of DNA nanogels at the atomic level can monitor the fabrication process and confirm the presence of targeting ligands.^11^ Advanced techniques can also give key insights into the stability of DNA nanostructures and evaluate their drug contents. Considering the cost of preparing DNA-based nanostructures, techniques that require little sample concentration and volume are warranted.

Overall, nanosized bioparticle analysis plays a crucial role in many areas of biomedical research and has significant potential to impact human health and well-being.^3,11^ However, the routine characterization, identification, and quantification of bioparticles is still challenging due to the lack of quick, sensitive, reproducible, and low-cost methods that use low sample concentrations and volumes.^1,8^ Among approaches, nanoparticle tracking analysis (NTA) is quick and useful to find out the size of the nanoparticles by tracking its Brownian motion and measure the zeta potential to determine the surface charge and stability in a liquid suspension.^12^ Electron microscopy is useful to provide high-resolution images and insights into the shape, size, morphology, and chemical compositions of nanocarriers.^6^ Other methods like enzyme-linked immunosorbent assay (ELISA) detect target specific proteins by immobilizing antibodies. Similarly, flow cytometry uses fluorescent labelled antibodies to quantify bioparticles in a liquid suspension. Although powerful, these methods do not provide the chemical fingerprint of the nanocarriers and are often low throughput and labour-intensive.^13^ Consequently, there is an increasing need for advanced techniques that can be used to analyse bioparticles with greater sophistication.^14^

Raman spectroscopy is an effective analytical tool for identification of molecules through chemical finger prints.^15^ However, Raman scattering is usually very weak, with the small cross-section of nanosized bioparticles additionally providing low intensity compared to the autofluorescence spectroscopy.^16^ Raman scattering from bioparticles such as EVs and liposomes are therefore typically performed using high laser power and long spectrum acquisition time.^17^ An alternative option is to use surface enhanced Raman spectroscopy (SERS), a technique that offers highly enhanced spectrum intensity at a lower laser power within a shorter acquisition times.^18^ SERS allows for real-time characterization without additional extended sample preparation requirements, enabling detection of molecules at trace levels.^19,20^ SERS has been utilized for bioparticle analysis in a variety of applications including drug delivery, disease diagnostics and monitoring.^21,22^ Also, the identification and detection of DNA and RNA sequences or bonds within crosslinked DNA nanostructures, changes and mutations have been explored using SERS.^23^ In SERS, metallic nanoparticles, such as gold (Au) or silver (Ag) can trap the incident laser light at minimal volume (a so-called “hotspot”) through the localized surface plasmon resonance (LSPR) effect.^19^ When an analyte is placed at the vicinity of the hotspot, its scattered Raman signal will be enhanced due to the high local electric field. As it is a near-field optical effect, having the analyte within a range of 10s of nm to the hotspot is essential. To cater to this need, SERS signal from a nanosized liposome is usually obtained by encapsulating plasmonic nanoparticles such as gold and silver nanoparticles within the liposomes itself.^24^ This method, albeit good for the characterization, poses challenges such as changes in membrane morphology, alteration of membrane fluidity, it can lead to membrane disruption and may trigger toxicity as well as in vivo accumulation.^25,26^

As opposed to attaching metal nanoparticles to the analyte, another route is to use a SERS substrate with defined hot spots at the surface using nanofabrication techniques, such as lithography, physical and chemical deposition methods, etc. The main challenge in this approach is the high Brownian motion of nanosized bio-particles in the solution, which alters the distance between the substrate and the nanoparticle influencing both SERS signal intensity and the repeatability.^27^ Several routes proposed to avoid high Brownian motion of nanosized bio-particles can be broadly classified in two approaches. The first approach is to either dry or stick the nanosized bio-particles on top of the surface. The challenge associated with this approach is that there might be protein breakdown and conformational changes of the fragile bio-particles in the dried samples which compromise the SERS signal.^28^ Characterization of biological samples in dried state leads to change in morphology,, rupture of lipid membranes, can induce pores, and cracks into the membrane of the biospecimen.^28^

A second label-free approach is to trap the nanosized bioparticles in liquid media to reduce the Brownian motion and to bring them in close proximity to the SERS substrate. Here, researchers have used conventional laser tweezers,^25^ microfluidic trapping,^29^ physical trapping via nano-structured surfaces (e.g., nano-hole)^30^ and electrophoretic trapping.^31^ Among the aforementioned methods, trapping using nano-structured surfaces does not add any additional instrumentation and is a straightforward method for high-speed characterization. Few literature references reported the SERS characterization of bioparticles using a nanostructure surface trap.^30,32^ Based on the analyte integrity of this approach, it is desirable to further develop such topologies that can trap and identify distinct chemical composition and Raman spectral data of a diverse nano-biocarriers.

Here, we optimized the design and the fabrication process of a flexible SERS substrate by means of a honeycomb-like nano-bowl morphology. The proposed honeycomb inspired nano-bowl SERS substrate provides a large electric field enhancement and enables localization by passively overcoming the Brownian motion of nanocarriers, thus improving the reproducibility of the SERS signal. The topology was fabricated out of a polydimethylsiloxane (PDMS) film through a self-assembled polystyrene beads (PS) monolayer template. The proposed design enables acquisition of SERS data from liposomes, DNA nanogels, and EVs in the liquid state to maintain their biological value. The nano-bowl structures offer more area and hotspot sites as compared to a flat surface, enhancing the Raman activity.

The nano-bowl SERS substrate was employed to obtain spectra from liposomes with different lipids such as phosphatidylcholine (PC) and phosphatidylethanolamine (PE) and from liposomes with their membranes modified with cholesterol. The spectral signatures of cholesterol were identified for both the lipoids with cholesterol samples. Surface modification with polyethylene glycol (PEG) further revealed improved SERS activity featuring characteristics bands of PE lipid membrane and additional Raman peak of PEG. The nano-bowl SERS substrate allows distinction between both the surface modified liposomes and the membrane modification of the lipid bilayer.

Next, we demonstrated the utility of the SERS substrate for label-free characterization of EVs. Distinct EVs subtypes derived from three different malignant leukaemia cells (HAP1, HAP1-F3KO, and THP1), were analyzed using the developed method to explore molecular signatures of EVs and the associated cell of origin. Thus, the EV subtypes derived from the mentioned cell lines contain a variety of biomolecules from the host cell, which helps finding out the role of intercellular communication and disease diagnosis. The SERS performance of all nanocarriers was evaluated by comparing their standard Raman spectra with SERS to estimate the Raman enhancement and determine the suitability of this technique to analyse such nanocarriers.

DNA nanogels (60 nm in diameter) were further used to further investigate the efficiency of our proposed nano-bowl substrate on sub-100 nm bioparticles. The SERS spectra of the cross-linked nanoparticles revealed the presence of compositional bases and ribose sugars, confirming the ability of SERS to directly characterize DNA nanogels in liquid without a typical encapsulation with metal nanoparticles.^10^ Fig.1 shows the schematic representation of model of nano-biocarriers used in this study.

**Fig. 1.**
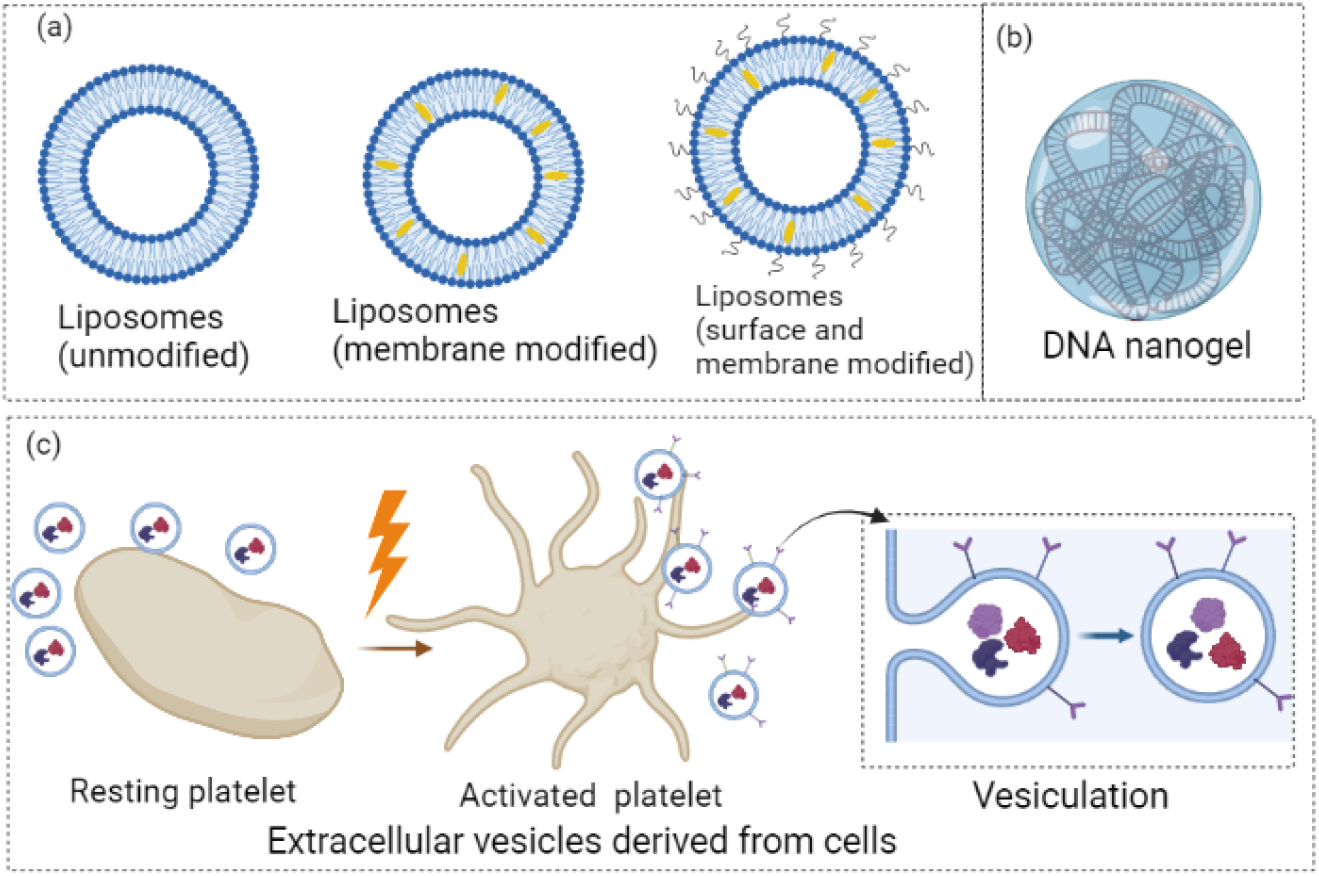
Schematic representation of nanosized bioparticles used in this work. (a) The membrane bilayer of liposome, attachment of cholesterol in the lipid bilayer, and surface and membrane modified liposomes (b) model of the DNA nanogel, (c) Schematic diagram of platelets and its activation that results the vesiculation of EVs.

## Methodology

### Materials

Si wafer, Polystyrene (PS, 190 nm, 500 nm, 1.04 µM diameter (10%)) beads from bang laboratory, Deionised (DI) water, Sodium dodecyl sulfate (SDS, > 99%), SYLGARD®184 is a silicone based elastomeric kit, Dichloromethane (>99.8%), Ag target.

### Substrate fabrication details

The nano-bowl structure was fabricated using the three different monolayer polystyrene (PS) beads stamps of 190 nm, 500 nm, and 1 µM diameter, respectively. The monolayer of PS beads was achieved using monolayer assembly at the air-water interface.^33^ The beads suspension in deionized (DI) water having 5% concentration was spin coated on plasma treated clean silicon (Si) wafer. The spin coating enables a partial monolayer of beads, and the coated Si wafer was slowly immersed in a petri dish containing DI water. This process enables a transfer of beads layer from Si wafer to the water. The surface tension of water was changed by adding 1.6 ml of SDS solution into 40 ml of DI water. Once SDS was added, a monolayer bead floating on the air-water interface was immediately observed and it was further adsorbed in another fresh Si wafer. For better adsorption of beads on Si wafer, the coated Si wafer was kept at 110° C for 20 mins. Then a bubble free PDMS solution was spin-coated at 800 rpm for 20 sec on the PS bead template for soft lithography. The PDMS solution was kept at 70°C at hotplate for 2 hours. Once it is cured, the PDMS was peeled off, hence a pattern of honeycomb-like nano-bowl structure on PDMS was formed (Fig. 2). It was further washed using dichloromethane and DI water to remove any bead residue from the surface and dried by nitrogen gun. Next the nano-bowl template was coated with silver (Ag) for varying thickness of 40 nm, 70 nm, and 100 nm, respectively, using a sputtering unit.

**Fig. 2.**
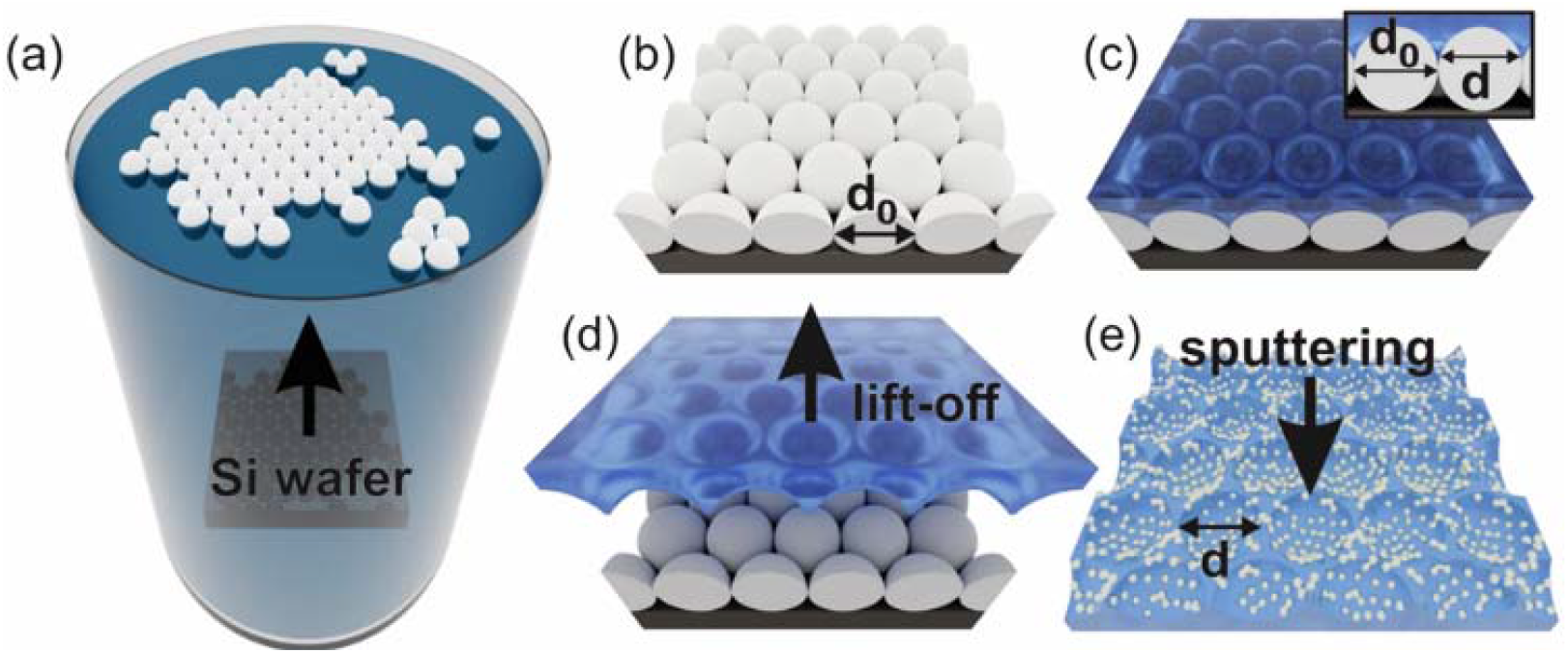
Schematic representation of fabrication of Ag coated PDMS nano-bowl SERS substrate. The monolayer PS beads template was formed at the air-water interface, which was further adsorbed slowly on a clean Si wafer. The PS monolayer on Si wafer was covered with PDMS, then cured, and peeled off. Thus, nano-bowl geometry on PDMS was formed. The nano-bowl was further Ag coated using a sputtering unit.

### Determination of plasmonic enhancement using 3D FDTD simulation

The 3D finite difference time domain (FDTD) simulations using Lumerical software were performed to investigate the distribution of hotspots with an enhanced electric field. The simulation model was constructed using a simplified model of the nano-bowl structure as an inverted hemisphere. The sphere diameter was varied for three different diameters of 90 nm, 300 nm, 550 nm, respectively, considering the structure parameters, such as diameter and spacing obtained experimentally and extracted using FESEM images. Although not fabricated, the simulation of 700 nm diameter nano-bowl was also performed in addition to 3 other diameters (90, 300 and 550 nm) to determine the best nanostructure for the maximum SERS activity. The thickness of Ag coating was set to be 70 nm for all geometry, due to optimised enhancement activity considering 40nm, 70nm, and 100nm Ag coated SERS film, shown in supplementary figure S.F. 1. The refractive index of PDMS was set at 1.43, and the refractive index of Ag was taken from the Palik data set. The simulation geometry was considered for a single bowl to restrict the simulation memory requirements. The incident laser profile was chosen to be Gaussian, with an excitation wavelength of 532 nm, a bandwidth of ±5 nm and is propagating in the –Z direction. The simulation domain was set for perfectly matched layer (PML) boundary conditions. The mesh size was kept auto-nonuniform, and the override region closing the entire nano-bowl was set as 3 nm for all dimensions. The frequency-domain field profile monitors were utilized to check the local electric field map at various planes of the nanostructure.

### Bio-Sample preparation

#### Synthesis and characterization of liposomes

Liposomal formulations were synthesized via the thin-film hydration method (1). Pure soy phosphatidylcholine, Lipoid S100 (SPC) or SPC containing cholesterol (CHOL) solutions were prepared by dissolving the lipids in methanol to achieve a percentage molar ratio of 100 (SPC) and 80:20 (SPC/CHOL). For PE containing liposomes, Lipoid E PE (EPE), cholesteryl hemisuccinate (CHEMS) and Lipoid phosphatidylethalonamine 16:0/16:0 PEG 5000 (PEG 5000) was dissolved in 50:50 solution of chloroform/methanol with or without cholesterol to achieve final percentage molar ratios of 54/41/5 (EPE/CHEMS/PEG 5000) or 44/31/20/5 (EPE/CHEMS/CHOL/PEG 5000) respectively. The solutions were placed under a vacuum at 45 ^°^C for 1 h. Thereafter, the film was rehydrated with phosphate buffered saline (PBS, pH 7.4) to achieve final theoretical concentrations of 10 mg/mL (SPC containing liposomes) or 8.5-8.9 mg/mL (EPE containing liposomes) of the lipids. The solutions were then extruded 6 x through each 800 nm, 400 nm, 200 nm, and 100 nm polycarbonate membrane. The liposomes were stored at 4 °C until used. The hydrodynamic diameter of the liposomal formulations was determined using dynamic light scattering (DLS) with the NanoZS Zetasizer (Malvern). The size measurement chart is provided in the supplementary section: 2 of the supporting information files.

### Synthesis and characterization of DNA nanoparticles

DNA nanoparticles were prepared as previously described with varying molar ratios of the nanostructures.^34,35^ Three DNA nanostructures namely, Y-SAB, Y-SAF and L-SAC were prepared by dissolving the oligonucleotides in 1x encapsulation buffer (5mM Tris-HCL, 1mM EDTA, 10mM MgCl_2,_ and 10mM NaCl) and annealed via thermal hybridization for 2.5 h. Equal volumes of the Y-SAF, Y-SAB and L-SAC monomers were then mixed to achieve molar ratios of 4/1/6.5 or 16/4/26 and hybridized at 95 °C. The solutions were then rapidly cooled to 4 °C and further incubated for 3.5 h. The hydrodynamic diameter of the nanoparticles was determined with a low-volume quartz cuvette via dynamic light scattering (DLS) with the NanoZS Zetasizer instrument, Malvern.

### Isolation of extracellular vesicles

Haploid human cell line HAP1 and its derivative TF-knock out cell line (KO F3) were cultured in IMDM supplemented with 10% FCS. Extracellular vesicles were isolated from haploid human cells (HAP1) and its tissue factor (TF)-knockout (KO) derivative HAP1-F3KO cells (Horizon Discovery Ltd.) and the THP1 monocytic cell line (American Type Culture Collection, ATCC). HAP1 and HAP1-F3KO cells were cultured in IMDM supplemented with 10% filtered fetal calf serum FCS; THP1 cells were cultured in RPMI supplemented with 5% exosome free FCS. EVs were isolated from the culture media of HAP1 and HAP1-F3KO by sequential centrifugation and resuspended in 20 mM HEPES/150 mM NaCl buffer. Size exclusion chromatography was used to isolated EVs from THP-1 and stored in PBS (See supplementary 2 of supporting information for more details). Isolated EVs were stored at -80°C until use.

### SERS spectrum acquisition

The fabricated SERS chip was exposed to nitrogen plasma to increase surface adhesion. Next, 2µL of sample solution was placed inside a rectangular PDMS chamber (1.5 micron thick), placed on the plasma treated SERS substrate. Then, the sample fluid was sealed using a glass coverslip (18mm thick) on top of it. The schematic representation of sample preparation protocol for SERS measurement is shown in supplementary Figure S.F. 2. The adopted strategy ensures that the material in solution does not dry out, preserving its biological value. All SERS spectra were acquired using Renishaw In-Via Micro Raman spectrometer (100X, 0.85 NA). All SERS measurements were performed using a 532 nm excitation laser, 10 milliwatt laser power, and 5 sec integration time, respectively. The collected spectra were baseline corrected, and background subtracted to perform further analysis. The collection of SERS spectra of samples prepared with the same methodology was also performed using a 785 nm laser. In that case, the SERS spectra of the sample overlapped with a broad fluorescence background at the crucial wavelength range (1200-1400-cm^-1^) coming from the coverslip, making it noisy, and hard to interpret.

## Results and discussions

### Optimization of SERS substrate

The structural parameters of nano-bowl SERS chip were optimized inspecting two important parameters: A) the nano-bowl structure, the diameter, and the height to be larger than the bioparticles, such as, liposomes (150 nm), DNA (60 nm), and EVs (100 nm)^32^ to entirely trap the particles; and B) the morphology of nano-bowl to exhibit enhanced plasmonic activity considering 532 nm excitation laser.

The smallest size of the bioparticles used in this work was 60 nm for DNA nanogel and the largest was for liposomes around 150 nm. Thus, we used the polystyrene beads of diameter 190 nm, 500 nm, and 1 µm to make the PDMS stamp. The morphology of structured PDMS surface cured from polystyrene monolayer beads template of varying diameters (190 nm, 500 nm, and 1 µm, respectively) without Ag coating is shown in Fig. 3 (a-c), respectively. The average bowl diameters were calculated to be 90 nm for 190 nm template, 300 nm for 500 nm template, and 550 nm for 1 µm template, respectively using ImageJ software. The obtained bowl diameter range between 90-550 nm was sufficient to trap DNA nanogels, EVs and liposomes.

**Fig. 3.**
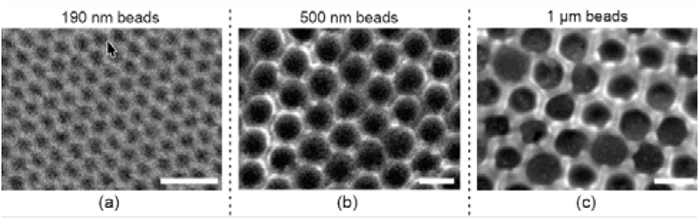
Field emission scanning electron microscopy (FESEM) image of nano-bowl structured PDMS film cured from a PS template of (a) 190 nm, (b) 500 nm, and (c) 1 µm. Scale bar: 500 nm.

Next, we investigated the amount of the plasmonic enhancement offered by the opted nano-bowl geometry as shown in Fig. 4 (a-e). The schematic diagram FDTD simulation set-up was shown in Fig. 4 (a). This was inspected using the 3D FDTD simulations considering Ag coated nano-bowl structures for four different diameters of 90 nm, 300 nm, 550 nm, and 700 nm respectively. The choice of the diameter was to mimic similar diameters obtained experimentally i.e. 90 nm, 300 nm, and 550 nm obtained using polystyrene beads of diameter 300 nm, 500nm and 1 µm, respectively. We also simulated a nano-bowl of 700 nm diameter, although not fabricated to investigate the plasmonic enhancement.

**Fig. 4.**
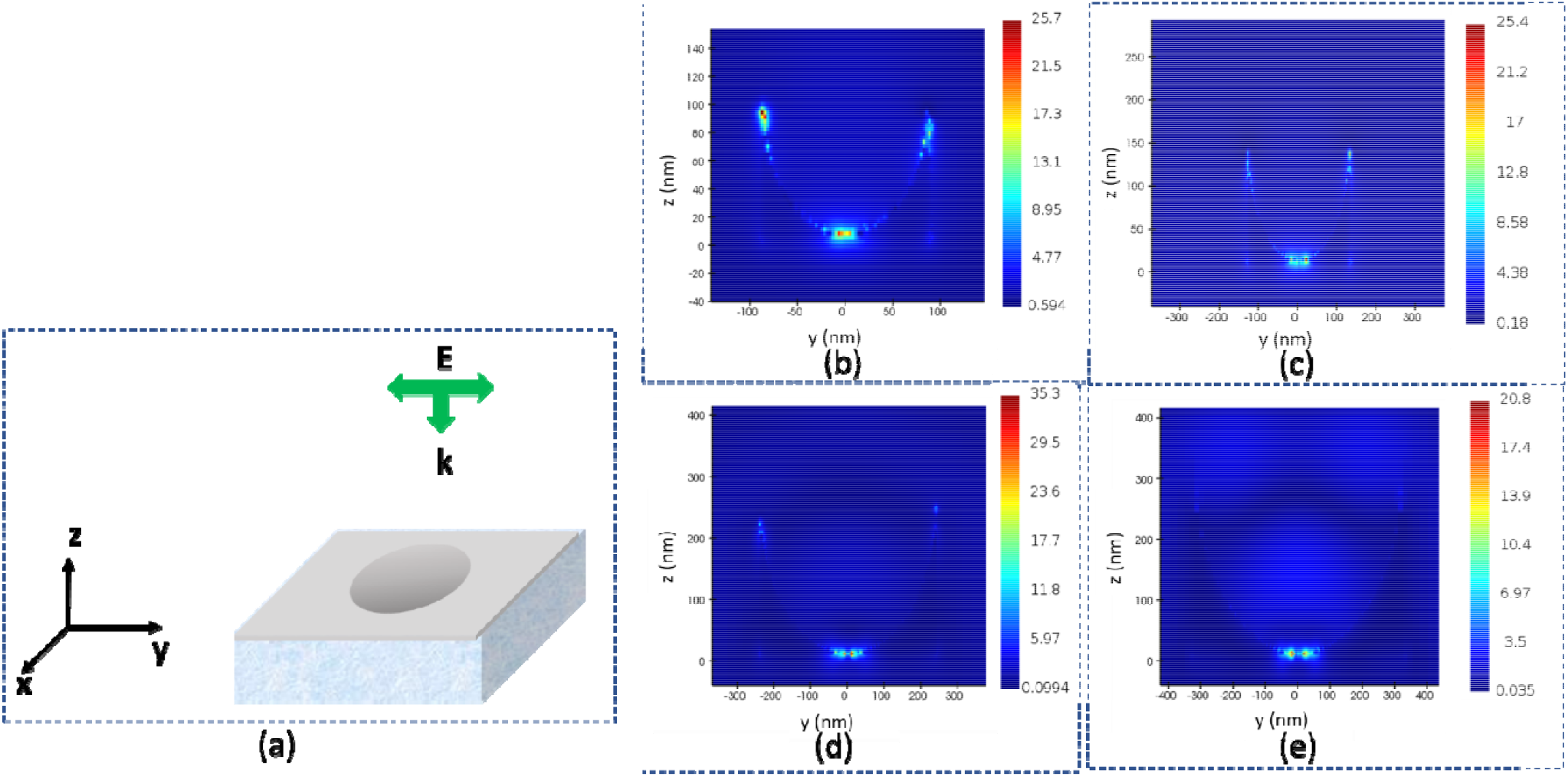
(a) Schematic of FDTD simulation set-up. (b) Mapping of Local electric field along YZ plane for fabricated nano-bowl geometry having diameters of (a) 90 nm, (b) 300 nm, (b) 550 nm, and (c) 700 nm. The hotspots occurred on the curved surface. The colour bars indicate the ratio of local E field to the incident E field.

Fig. 4 (b-e) revealed the local field map along the YZ plane for different nano-bowls’ diameters. The simulation results displaying the nano-bowl of 550 nm diameter prescribed best enhancement for 532 nm laser wavelength, considering all other simulation parameters constant. The reason behind the maximum SERS enhancement for the bowl structure is the ability to produce enhanced electric field due to curvature induced focusing of electromagnetic field because of the curved surface.^36^ The curve structure, acting as a lens, focuses the laser light spot tightly creating higher intensity of hotspots compared to the flat surface.^37^ The shape anisotropy of the Ag-PDMS film also induces higher density of hotspots compared to a flat film. The increased plasmonic activity for the curved surface is due to the surface plasmon polariton (SPP) waves generated at the metal dielectric interface due to the continuous metallic film. The SPP wave travelling from edge to bottom of the nano-bowl surface gets a large area for enhanced electric field. The coupling between SPPs is strongly affected by the bending of surface, since the travelling of SPP is guided through the surface structure and leading to a low-loss transmission of plasmonic signal.^36,37^ The optimisation is applied to the simplified model of flat Ag film compared to the real geometry, where we can expect some discontinuity in thee Ag structure because of the curvature of the surface. The discontinuous Ag will exhibit LSPR for the strong enhancement in the Raman signal.

The chosen nano-bowl of a diameter of 550 nm were further coated with 40 nm, 70 nm, and 100 nm Ag to experimentally investigate the best SERS activity. We found similar enhancements for three different Ag coating thicknesses, considering the SERS spectra of 1 µM R6G, shown in supplementary Figure S.F.1. The reason behind similar enhancement is due to the minor thickness variation of Ag. If the thickness is too small, the number of hotspots will be less. Due to the decreased number of hotspots, the enhancement factor would decrease. If the Ag film is too thick, the surface will be flat, and curvature will be reduced, lowering the enhancement factor. Therefore, when the thickness of the Ag film is within the optimal range, the SERS enhancement factor is at its maximum, and the SERS intensity for R6G was not affected by the minor changes in the film thickness. Therefore, choosing the optimal Ag coating range for the fabricated nano-bowl surface, the 70 nm Ag coated PDMS SERS film was used for all measurements.

Another principal factor of SERS substrate quality is the ability to reproduce the enhanced Raman spectrum over the entire surface of a particular SERS substrate, choosing random locations and substrates obtained from different fabrication batches. The optimized SERS substrate produced reproducible Raman results for 1 µM rhodamine 6g (R6G) considering seven random spots on a single substrate, as shown in Fig. 5 (a). The spectra reproducibility was further investigated by considering three fabrication batches with R6G as a probe molecule. The spectra are highly reproducible; peaks occurred at the same wave shift value, and the intensity variation was calculated to be less than 5%. Further the substrate can identify the peaks up to 10^−15^ M concentration of R6G (Fig. 5 (b)). The peaks at 608 cm^-1^ represent the bending of C-C-C ring, peak at 773, 1126 cm^-1^ for in plane bending of C-H group, 1309 cm^-1^ for in plane bending mode. The peaks at 1361, 1505, 1574, and 1649 cm^-1^ are assigned for aromatic C-C stretching.^38^

**Fig. 5.**
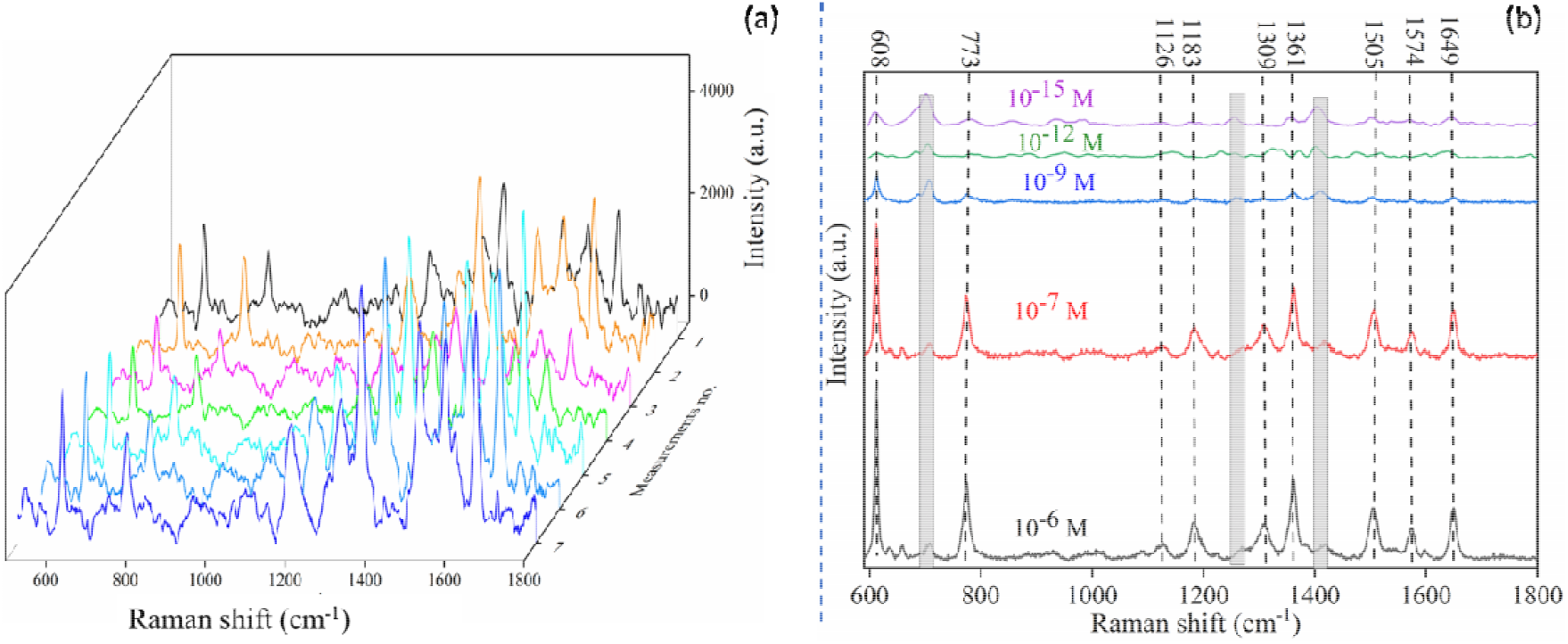
(a) Reproducibility of SERS substrate. Acquisition of 1 µM R6G SERS spectra over 7 random spots. The peaks are reproducible and intensity variation is less than 5%. (b) SERS spectra of R6G for 10^−6^M -10^−15^ M concentration. The nano-bowl SERs substrate detects enhanced and distinct peaks of R6G up to 10^−15^ M concentration. The shaded peaks indicate the Raman spectrum of PDMS substrate.

### Analysis of trapping activity by nano-bowl PDMS structure

The ability to hold polystyrene nanoparticles inside the nano-bowl geometry was studied using the FESEM. First, the substrate cured from 1 µm polystyrene template was plasma-treated to improve surface wettability. The plasma-treated bowl has a wall surface barrier that exert friction against the smooth flow of the fluid. The increased surface adhesion will also contribute to the additional frictional forces. Thus, the nanobeads with a diameter less than the curvature falls inside the bowl, cannot overcome the frictional force, and will be eventually localized. To facilitate the flow of the liquid, the surface was tilted at an angle so the liquid could move from one end to the other. This technique avoids the aggregation of beads at random spots. Fig. 5 (a) shows the SEM image of 250 nm polystyrene nanobeads captured inside the nano-bowl, demonstrating the localization of individual nanobeads inside the nano-bowl.

To evaluate the effectiveness of nanoparticles trapping, we tracked the motion of liposomes using a bright-field video microscope (60X, 1.2 NA water immersion objective lens). The captured videos were uploaded and shared in the supplementary file of supplementary section 1. Screenshots of extracted frames of EPE liposomes placed on nano-bowl structured and flat PDMS are provided in supplementary Figure S.F.3. The particles’ coordinates (X, Y) were extracted using ImageJ software, and the trajectories of un-trapped, entirely trapped, and partially trapped lipids were plotted, shown in Fig. 6 (b-e). Motion tracking allowed us to classify the motion of liposomes into three categories:

**Fig. 6.**
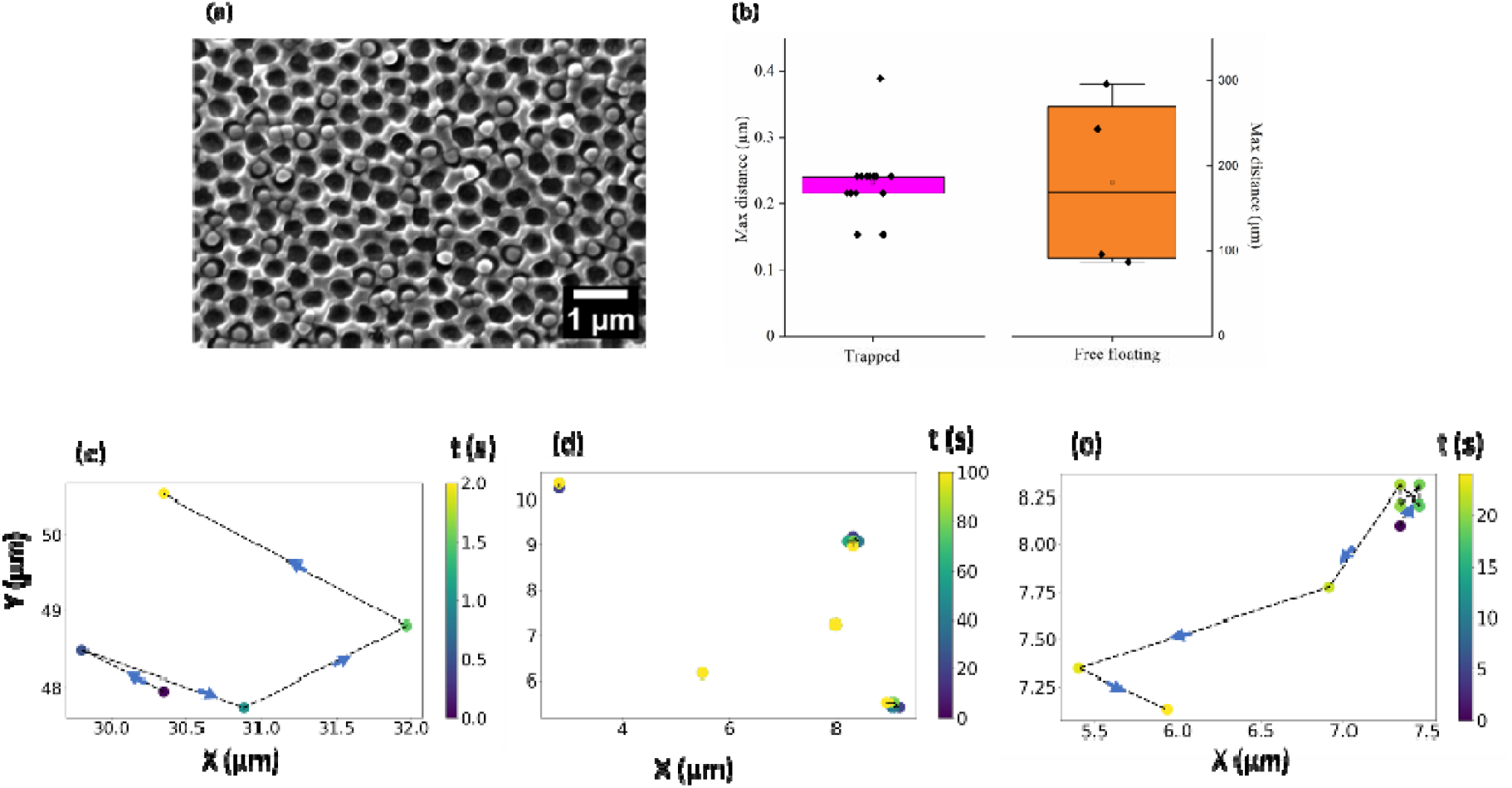
(a) SEM image of trapped polystyrene beads (250 nm diameter beads) inside the nano-bowl, cured from 1µm PS template. (b) Shows the maximum displacement of trapped liposome for up to 100 s while the free-floating liposomes were only tracked for up to 4 s due to high Brownian motion. (c) Dynamic track plot of a free-floating liposomes on flat PDMS, (d, e) show trapped and partially trapped liposomes on the nano-bowl shaped PDMS respectively. The plot implies that the trapped liposomes are localized, easy to focus, and while the free and partially trapped liposomes exhibit arbitrary diffusive motion over time. The tracking in (c-e) was only performed in XY plane, the free-floating liposomes had a significant motion along the Z-axis as shown in Supplementary Movie. S.M. 1 and S.M. 2 of supplementary section 4.

1. Liposomes free-floating on a flat PDMS film displayed random Brownian motion with rapid in-and-out of focus plane, making them very difficult to track even for a few seconds, see Fig. 6 (c).
2. Liposomes that were trapped by the honeycomb-like nano-bowl surfaces. Although liposomes tried to escape from the trap, its coordinates remained the same over longer period of time several minutes making them highly localized, see Fig. 6 (d).
3. Liposomes that were partially trapped to another liposomes. This occurred when a free-floating liposome attached to another trapped liposome, becoming partially attached to the nano-bowl surface. Since it continuously moved, it escaped the trap after a short internal, for example after 19 seconds as shown in Fig. 6 (e).

### SERS Analysis of Liposomes

The fabricated SERS substrates were utilized to examine the real-time characterization ability of a diverse set of liposomes. We examined two aspects for characterization, suitable for therapeutic application. First to investigate if nano-bowl SERS substrate can identify two differently composed liposomes and secondly if it can show spectral features that are unique to membranes, such as membrane modification. Hence, the SERS spectra of two different phospholipid compounds of PC and PE, and their membrane structure modified with and without cholesterol were examined, as shown in Fig. 7.

**Fig. 7.**
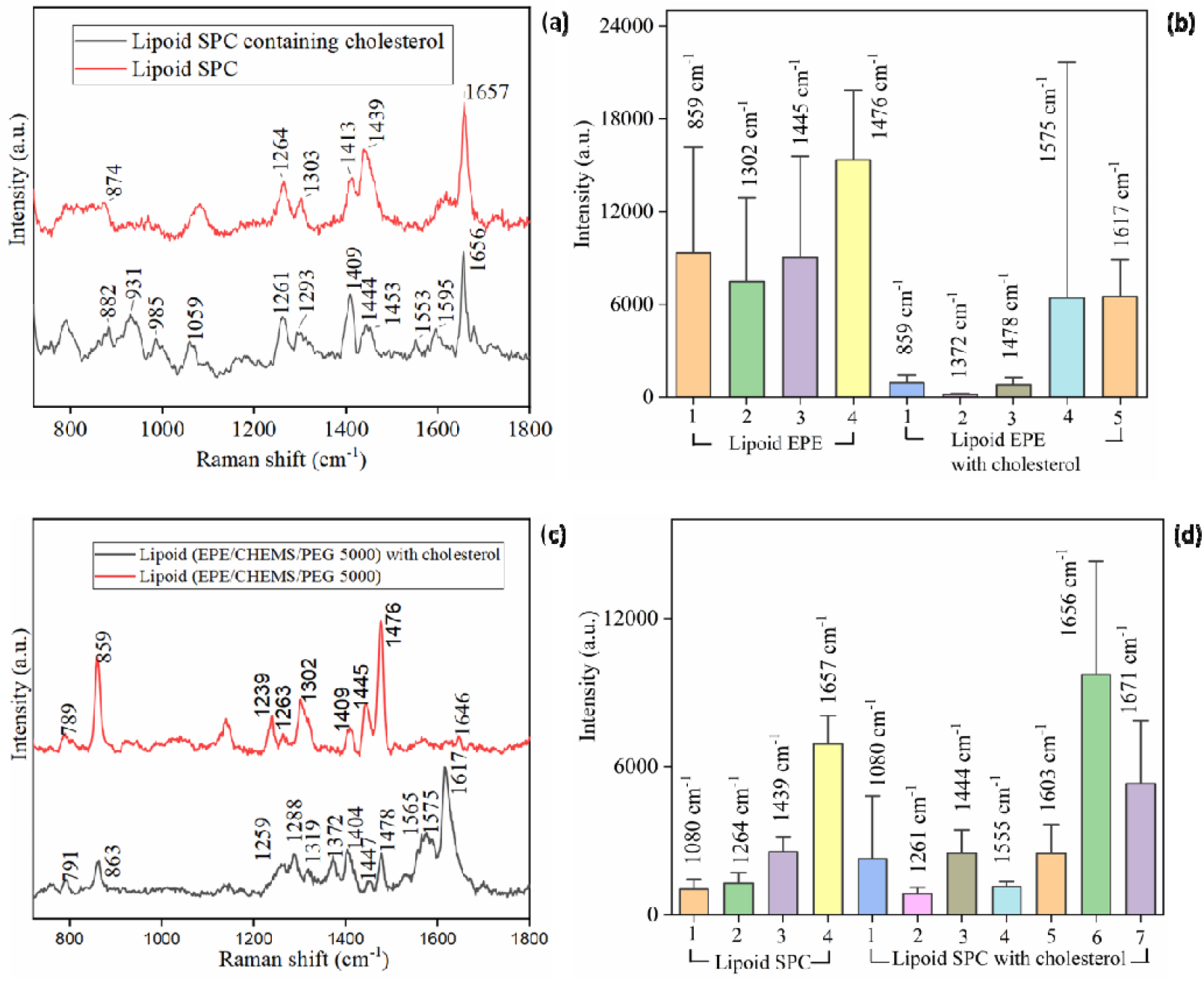
SERS spectra of liposomes (a) SPC with and without cholesterol, (c) EPE with and without cholesterol. The membrane structure gets shifted and additional Raman peaks occur due to the attachment of cholesterol into the lipid membrane. Average distribution of intensity of the Raman shifts for 15 independent measurements of (b) SPC liposomes with and without cholesterol and (d) EPE with and without cholesterol. The specific peaks at 1555, 1671 cm^-1^ are occurred for SPC liposome with cholesterols. For EPE liposome with cholesterol, peaks at 1565, 1575, 1617 cm^-1^ are attributed to cholesterol.

In Fig. 7 (a), Lipid SPC exhibits Raman peak at 874, 1264, 1303, 1439, and 1657 cm^-1^ are assigned for lipid bands.^25^ The SPC containing cholesterol gives shifted lipid band peak at 882 cm^-1^. The peak at 1059 cm^-1^ corresponds to stretching of C-C carbonate chains. The peak 1303 cm^-1^ is shifted to 1293 cm^-1^ for SPC with cholesterol. The additional peaks at 1553 and 1595 cm^-1^, 1737 cm^-1^ for ester linkage of cholesterol. The shift at 1672 cm^-1^ originates from the C=C stretching vibrations for cholesterol.^39^

Fig. 7 (c) shows the Raman spectra of another type of lipid (EPE/CHEMS/PEG) and the EPE lipid with cholesterol in the membrane. Some additional Raman shift observed at 789 cm^-1^ arising from the ethanolamine group for phosphatidylethanolamine spectrum.^40^ The shift at 859 cm^-1^ indicates the vibration of choline N+(CH_3_)^3^ group of phosphatidylcholines. The peak at 1239 cm^-1^ for EPE/PEG lipid appears twisting of CH_2_ bonds for PEG molecule. Peak at 1302 cm^-1^ for twisting of CH_2_ of PE lipoid. A highly intense peak at 1476 cm^-1^ assigns bending of CH_2_ of PE lipid membrane. For EPE lipid with cholesterol, additional Raman peaks at 1372, and intense peak at 1617 cm^-1^ are observed. The additional peak at 1372 cm^-1^ characteristic band occurs due to CH_2_ twisting mode of cholesterol oleate.

Fig. 7 (b), and (d) show the bar chart of Raman intensity distribution over fifteen independent measurements on random locations, for all liposome samples. The plot shows an enhanced intensity at 1656 cm^-1^ for SPC with cholesterol. Additional cholesterol peak intensity mapped at 1555 cm^-1^ and 1671 cm^-1^ are also key characteristics of cholesterol presence. An enhanced intensity of 1059 cm^-1^ is observed for cholesterol added SPC lipid^42,43^. For EPE lipid with cholesterol, additional peaks at 1565, 1575, and 1617 cm-1 are specific Raman peak of cholesterol. The normal Raman spectra of both liposome samples were acquired, and the SERS spectrum reproducibility was investigated, shown in supplementary sections 5, and 6 with supplementary Figure SF 4 and section 4 with SF 6, respectively.

The surface modification of EPE lipid using PEG/CHEMS was found to exhibit improved SERS intensity, as shown in Fig. 7. In the supporting information file, a comparison of normal Raman and SERS intensity was provided in Supplementary Figure S.F. 4 to further support this fact. The improved SERS performance can be attributed to the surface charge modification achieved using CHEMS/PEG. PEG is commonly used for stabilizing metallic nanoparticles, and this improved surface adhesion can affect SERS measurements.^41^ Additionally, PEG-functionalized lipids exhibit better hydrophilicity and stability.^42^ Thus, the evolved measurement technique allows for the sensing of both surface and bilayer modifications simultaneously, providing a useful tool for real-time characterization.

### SERS Analysis of DNA

The formation of the crosslinked DNA nanogels were previously confirmed and their sizes characterized with DLS. As shown in Table 1, the DNA nanogels revealed sizes of 62 nm which agrees with previous reports providing a model for SERS analysis.^10,43^ In Fig. 8, the SERS spectrum of DNA reveals a peak characteristic band at 791 cm^-1^, assigned for poly C and poly T ring breathing mode. The peak at 930 cm^-1^ indicates the single cytosine (C), peak at 1018 cm-1 for PO_2_-stretching of backbone, 1084 for OPO symmetric stretching, 1323 cm^-1^ for Adenine (A), 1476 cm^-1^ for A, T, C, and the shift at 1553 cm^-1^ and 1596 cm^-1^ for adenine band, respectively.^43^ The spectrum reproducibility was checked over seven random spots and is shown in supplementary figure S.F. 6(a) of supplementary section 6. The distinct peak at 1726 cm-1 is assigned for (C=O) bond of dG. The SERS spectrum shows the structure related Raman bands, indicating the possibility of SERS-based label-free detection of DNA nanogel using the developed methodology.

**Table 1.**
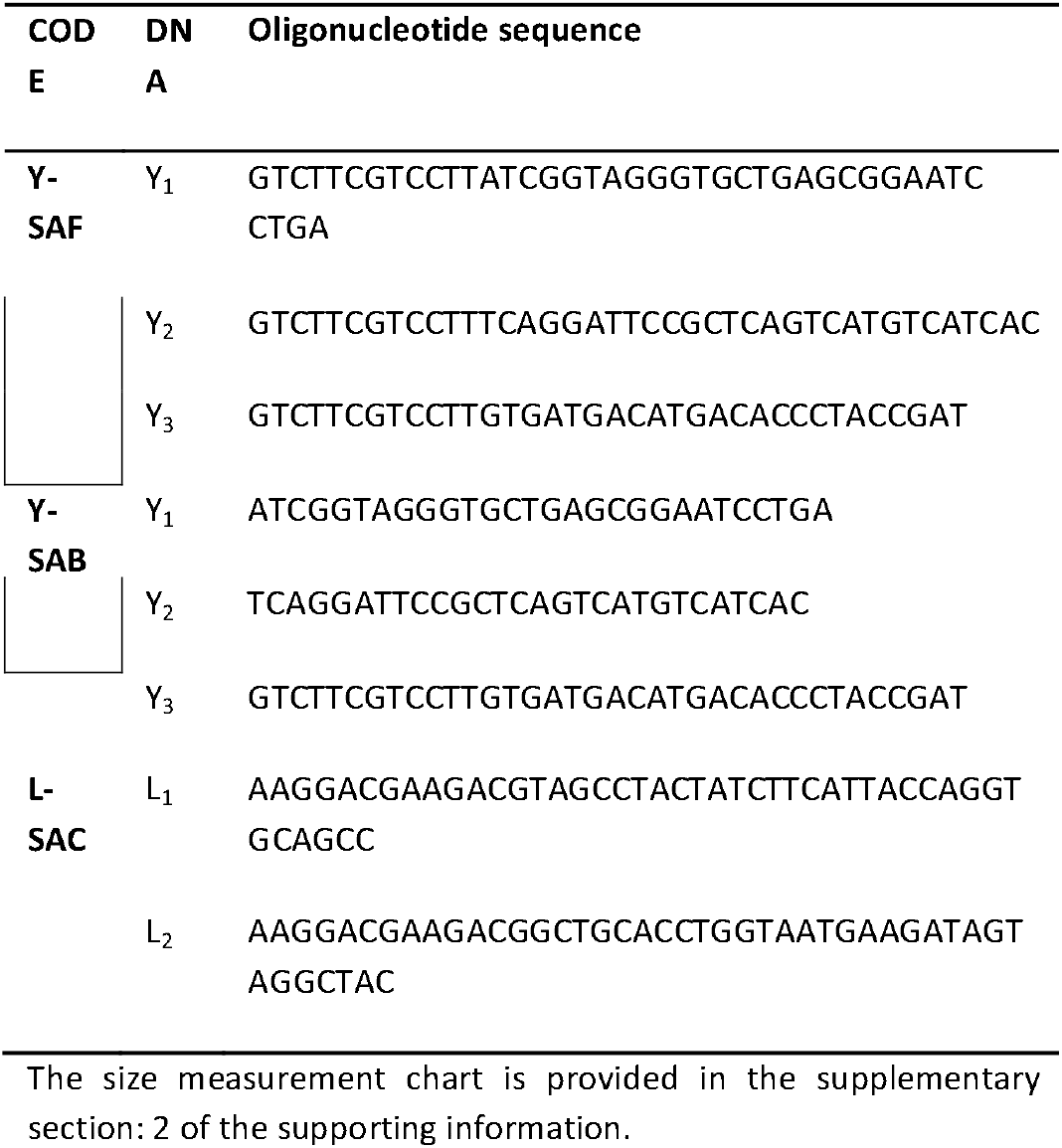
Oligonucleotide sequence of DNA nanogel sample

**Fig. 8.**
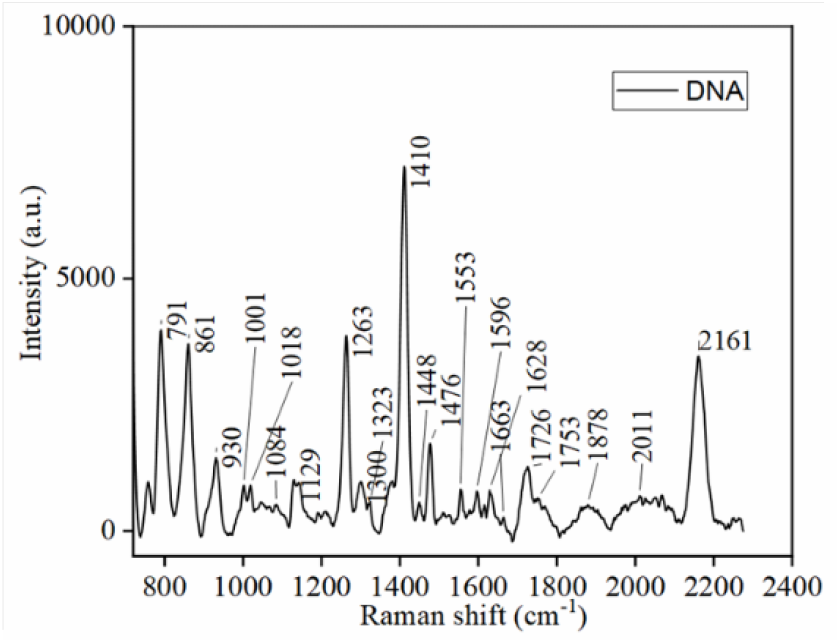
SERS spectrum of DNA nanogel indicating enhanced SERS peaks of DNA building blocks.

### Analysis of EVs

To further investigate the ability to distinguish and specify a diverse set of bio-nanocarriers, SERS spectra were acquired from EVs isolated from three distinct cell lines, THP-1, HAP1, and HAP1-F3KO, as shown in Fig. 9(a). The THP1 cell line is a monocytic cell line commonly used to model macrophage functions, intercellular communication, and signaling.^44^ The HAP1 cell has one copy of the chromosome, making it a valuable model for genetic research. HAP1 F3KO cells are genetically engineered HAP1 cells without the tissue factor (F3) gene, which has an essential role in blood coagulation. Thus, the EV subtypes derived from the mentioned cell lines contain a variety of biomolecules from the host cell, which helps find out the role of intercellular communication and disease diagnosis. The THP1-derived EVs show distinct spectral features, with a peak at 809 cm-1 attributed to nucleic acid. The SERS peaks are assigned at 1264 cm-1 for lipids, 1322 cm-1 for proteins, 1409 cm-1 for lipids, 1447 cm-1 for proteins and lipids, and 1620 cm-1 for proteins, respectively.^46^ Fig. 9 (b) represents intensity variation for three distinct peaks at 1264, 1409, and 1620 cm-1 considering 15 different measurements from THP1-derived EVs. HAP1-derived EVs show unique spectral features at 916 cm-1 attributed to nucleic acid, 1171 cm-1, 1354, 1566, and 1612 -cm-1 for protein. The SERS spectra of HAP1-F3KO-derived EVs exhibit Raman peaks at 788 cm-1, indicating the pyrimidine ring of nucleic acid. The shift at 880 cm-1 is assigned to proteins, 1264 cm-1 showing Amide III protein, 1140, and 1411 cm-1 for protease protein, respectively. The HAP1-F3KO derived EVs offer enhanced spectra at 1000 cm-1 for lipids, 1169 cm-1 for proteins, 1265 for lipids, 1352, 1411, 1515, and 1606 cm-1 for proteins, respectively.^47^ Fig. 9 (c-d) represents the intensity variation of intense peaks over 15 independent measurements considering HAP1 and HAP1-F3KO derived EVs, respectively.

**Fig. 9.**
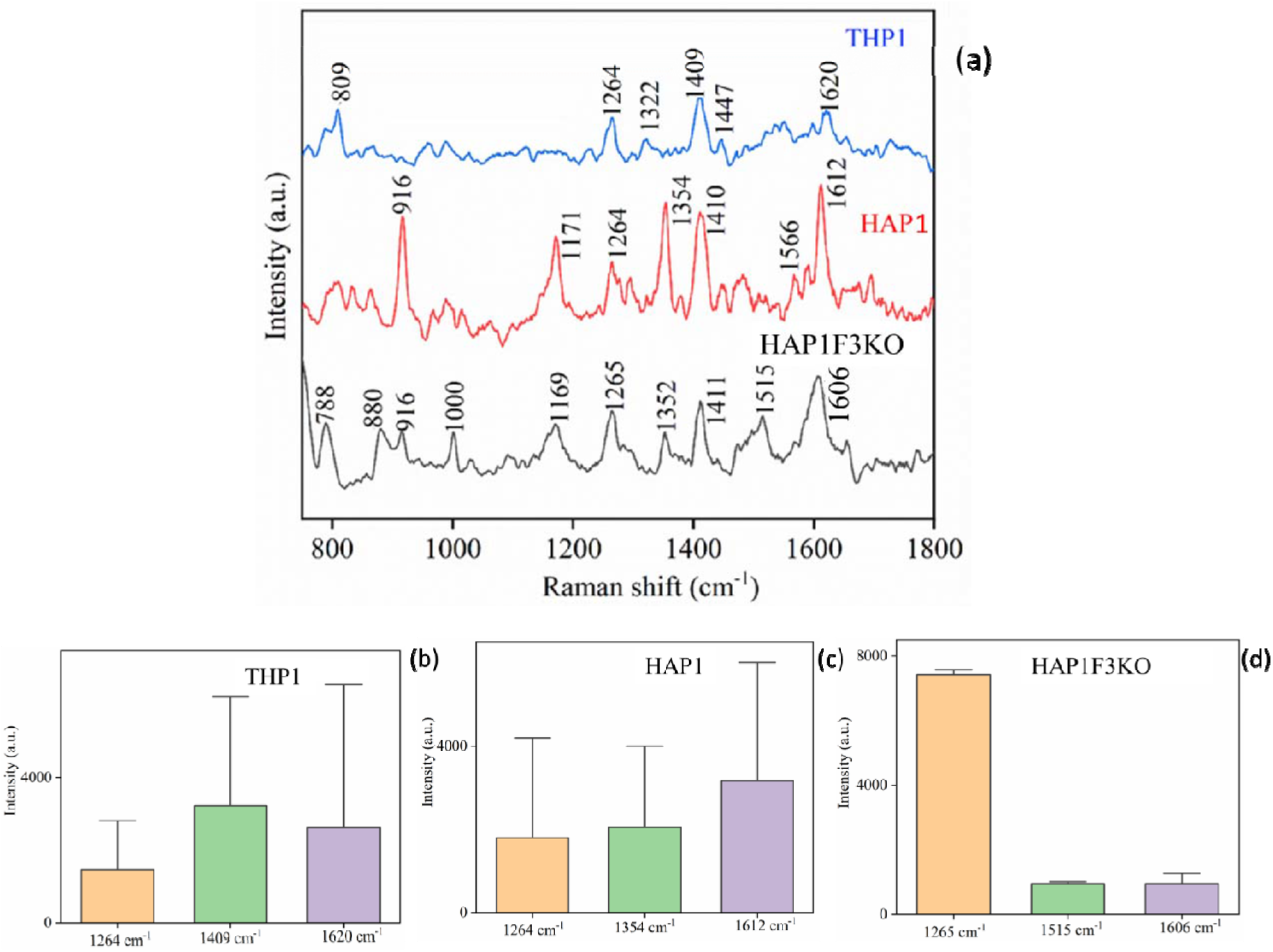
(a) Raman spectra of EVs from various sources isolated in HEPES buffer, respectively. (b-d) represents bar chart giving additional insights into the distribution of the dataset of corresponding Raman shifts for 15 independent measurements.

The peaks assigned for EVs are listed in the table below:^46,47^

| Raman shift | Assigned vibrational modes | Biomolecules |
| --- | --- | --- |
| 643-645 | Tyrosine | Proteins |
| 700-704 | cholesterol | Lipids |
| 717 | C-N<sup>+</sup> stretching | Phospholipids |
| 810-812 | Stretching (O-P-O) phosphodiester | Nucleic acid A type helix |
| 830-840 |  | Nucleic acid B type helix |
| 1000-1004 | Phenylalanine | Protein |
| 1050-1130 | C-C stretching | Lipids |
| 1125 |  |  |
| 1155-1160 | Stretching (C-C) of polyene | Carotenoids |
| 1175-1177 | Tyrosine | Protein |
| 1207-1210 | Phenylalanine (F<sub>3</sub>), Tyrosine (Y<sub>3</sub>) | '' |
| 1230-1305 | Amide III | '' |
| 1260-1270 | C-H<sub>2</sub> distortion in acyl chain | Lipid |
| 1295-1305 | '' | '' |
| 1300-1350 | Tryptophan | Protein |
| 1435-1465 | Distorsion C-H<sub>2</sub>, C-H<sub>3</sub> | Protein, Lipids |
| 1450-1490 | Guanine ring Purine A | Nucleic acids |
| 1515-1540 | Stretching polyene | Carotenoids |
| 1550-1555 | Tryptophan | Proteins |
| 1570-1580 | A, G ring | Nucleic acids |
| 1602-1604 | Ergosterol | Lipids |
| 1605-1607 | Phe (F1) | Proteins |
| 1615-1617 | Tyrosine (Y1) | '' |
| 1619-1621 | Tryptophan (W1) | '' |
| 1650-1670 | Acyl chain | Lipids |
| 1640-1700 | Amide I | Protein |
| 1670-1690 | C=O stretching [pyrimidine | Nucleic acids |
| 1720-1750 | C=O stretching in ester | Lipids |

A closer comparison of three EV subtypes revealed that the THP1-derived EVs do not show any distinct peak in the 1100-1200 cm^-1^ region dedicated to proteins and carotenoids. Further, the SERS spectra revealed that the peaks of THP1 EVs mainly occur due to functional groups of lipids and proteins. The shift at 1000 cm^-1^ for HAPF3KO is distinct and occurs due to protein structure. HAP and HAPF3KO show similar spectral features, indicating that their membrane composition is similar. Although there are similarities, HAP EVs show an enhanced peak at 916 cm^-1^, 1354 cm^-1^, 1566 cm^-1^, and 1580 cm^-1^, respectively, showing additional peaks of proteins and nucleic acids, which can be related to the state of the cells (viability, passage, proliferation etc.). The sharp peak at 1354 cm^-1^, and 1612 cm^-1^ for HAP1-isolated EVs is shifted to 1352 cm^-1^ and 1606 cm^-1^, respectively, with a decreased intensity for HAP1-F3KO derived EVs. Further, a clear distinction of spectral pattern for these two EV subtypes in the region (1500–1600 cm^-1^), where the protein peak at 1566 cm^-1^ and the peak due to the nucleic acid at 1580 cm^-1^ in HAP1 isolated EVs are not present in the HAP1 F3KO derived EVs.

In order to investigate the enhancement amount, a comparison was made between the Raman and SERS spectra of HAP1 and HAP1 F3KO isolated EVs, shown in Supplementary Figure S.F. 5. The Raman spectra of EVs derived from HAP1 and HAP1 F3KO showed similar spectral patterns, with only slight shifts in the Raman peaks observed in the 600-1000 cm^-1^ region. The spectral region of 1000-1600 cm^-1^ was hardly distinguishable between the two. In contrast, SERS spectra of the EVs showed distinguishable differences. This could be attributed to the fact that SERS is more sensitive to surface differences than Raman spectroscopy.

## Conclusion

We developed and validated a novel methodology to fabricate Ag-coated nano-bowl to trap and characterize nanosized liposomes, EVs and DNA-gel. The trapping was possible due to the surface topography of the novel nano-bowl pattern. The SERS signals of the membrane morphology, lipid compositions, and membrane modifications were detected for liposomes using the proposed nano-bowl design. The lipids comprising PEG exhibited a few additional peaks as compared to the lipids without PEG. The membranes of the SPC and EPE lipids had different vibrational peaks, indicating that it is possible to use nano-bowl SERS substrate to determine different membrane composition of liposomes (160 nm) in wet condition. Interestingly, incorporation of cholesterol in liposome membrane exhibited additional Raman features for distinction.

Further, the SERS methodology enabled identification of EV subtypes from diverse set of cell lines. Of note, although HAP1 and HAP1-F3KO cell lines are similar, the cells were cultured separately and the differences that are shown here between EVs from the two different lines may reflect the differences between the situation of cells. Although the SERS intensity count was modest, the spectra of EVs displayed additional vibrational peaks of a heterogeneous structure. Despite the additional vibrational peaks displayed in the SERS spectra of EVs, the SERS intensity count was low. We postulate that low enhancement is because the surface of EVs is electronegative, making it difficult for them to adhere well to negatively charged metals. This poor adhesion results in a low enhancement of peak intensity. So, in future, a surface modification can be developed to increase adhesion of EVs inside the trap. Finally, SERS methodology was able to identify the ribonucleic acids in DNA nanogel.

This investigation aims to develop proof of concept method to provide valuable details about the membrane modifications in liposomes, formation of DNA nanogel, and differentiation of EV subtypes without additional chemical reactions, or multiple characterization steps. In future the platform will be tested for the characterisation of these nanocarriers for targeted delivery.

## Author Contributions

Conceptualization, Fabrication of SERS substrate, Data curation, Formal Analysis, Methodology, Software, Writing – original draft: **Sathi Das;** Resources, Fabrication of SERS substrate, Discussions, Visualization, Writing, review & editing: **Jean-Claude Tinguely;** Preparation of Liposome, and DNA samples, Visualization, Writing, review & editing: **Sybil Akua Okyerewa Obuobi and Natasa Skalko-Basnet;** Isolation of EVs, Writing – review & editing: **Eduarda M. Guerreiro;** Supervision, Investigation, Writing, review & editing: **Omri Snir, Kanchan Saxena;** Supervision, Conceptualization, Funding acquisition, Investigation, Writing, review & editing: **B.S. Ahluwalia;** Supervision, Conceptualization, Funding acquisition, Investigation, Writing, review & editing: D. S. Mehta

## Acknowledgements

S. Das is thankful to Prime minister research fellowship (PMRF), Govt. of India. Research Council of Norway funded INTPART grant nanoSymBioSys (id. 309802). S. Das is extremely grateful to Prof. Matteo Chiesa, Renewable energy group, UiT Tromso, and FIST (DST Govt of India) UFO scheme of IIT Delhi for the Raman measurement facility. S. Das is thankful to Deanna L Wolfson for her support. S. Obuobi is thankful to the Tromsø Forsknings-Stiftelse (TFS) for the support through the TFS starting grant (20_SG_SO).

