## Supplementary material for "Honeycomb-inspired SERS nano-bowls for rapid capture and analysis of extracellular vesicles and liposomes in suspension": Suplementary

**Supplementary Section :1**

**Optimization Ag coating on patterned PDMS**

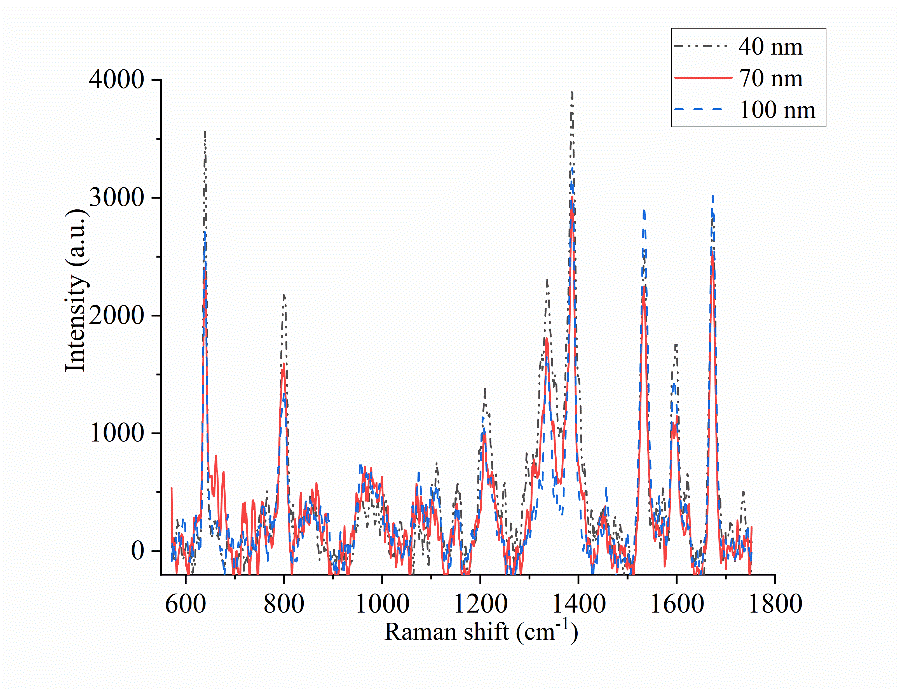

S.F.1 SERS spectra of 1 µM R6G on Ag coated PDMS SERS substrate, where the Ag coating thickness were 30nm, 70nm and 100 nm respectively.

**Supplementary Section :2**

**Table S1. Size measurement results:**

| No. | Formulation | Mean diameter (nm) | Polydispersity index (PDI) |
| --- | --- | --- | --- |
| 1 | SPC (100) | 141.9  142.1  143.0 | 0.076  0.085  0.074 |
| 2 | SPC/CHOL (80/20) | 136.8  137.0  135.6 | 0.052  0.081  0.09 |
| 3 | EPE/CHEMS/PEG 5000 (55/40/5) | 143.0  147.3  146.2 | 0.158  0.154  0.171 |
| 4 | EPE/CHEMS/CHOL/PEG 5000 (44/32/5/20) | 143  141.7  143.5 | 0.123  0.119  0.124 |
| 5 | DNA NP (16/4/26) | 62.6  61.1  63.2 | 0.417  0.402  0.427 |

**Isolation of extracellular vesicles**

**HAP and F3KO**

Isolation of EVs from HAP-1 and HAP1-F3KO cells was carried out by seeding 3 million cells in T-175 flasks in 40 ml IMDM+10% FCS. Cells were incubated overnight at 37°C, 5%CO_2_. Culture media was collected and discarded, and cells were washed with warm PBS. Forty ml IMDM + 10% exosome free FBS were added, and cells were incubated for 24 hours at 37°C, 5%CO_2_. Following incubation, the culture media was collected and EVs were isolated by sequential centrifugation. Culture media was initially centrifuged at 300xg for 5 min in a 5810 Eppendorf centrifuge, swing bucket rotor A-4-81. Supernatant was transferred to a fresh tube and centrifuged at 2.500xg for 10 min in a 5810R Eppendorf centrifuge, swing bucket rotor A-4-81. Pellet was discarded and the supernatant was loaded into 50 mL, Polycarbonate Bottle with Screw-On Cap (Beckman Coulter) and centrifuged for 20.000xg for 30 min, 4°C in an Avanti J-26 XP centrifuge (Beckman Coulter) equipped with JA-25.50 fixed angle rotor. EVs were resuspended in 900 µl of 20 mM HEPES, 150 mM NaCl buffer and stored at -80°C until use.

**THP-1- derived EVs**

Cells were cultured in 20 ml of RPMI+5% exosome free FCS in T-75 flasks, 37°C, 5%CO_2_. Fifteen ml of cell suspension were collected and centrifuged 250gx for 5 min. Supernatant was transferred to a fresh tube and centrifuged at 2500xg for 10 min to remove cell debris. Thirty ml of cell culture supernatant were pooled and concentrated in an Amicon-Ultra 15 Centrifugal filter units (Ultracel-50, Merck Millipore). The cell culture supernatant was loaded into the Ultracel-50 and centrifuged at 4,000rpm for 10 min in a Megafuge 1.0 (Heraeus Sepatech) centrifuge equipped with a swing bucket rotor BS4402/A. The concentrated cell culture media was collected to a fresh tube and the final volume adjusted to 1 ml by the addition of filtered PBS. Next, the concentrated cell culture media was loaded into a pre-washed 10 ml Sepharose CL-2B (GE Healthcare Bio-Sciences) size exclusion chromatography. Sample was allowed to enter the column matrix and PBS was added continuously. The eluate was collected in 15 sequential fractions of 0.5 ml each. Protein quantification was used to identify the EV-rich fractions, which were then pooled and stored at -80°C until use.

**Supplementary Section :3**

**SERS spectrum acquisition of biomolecules**

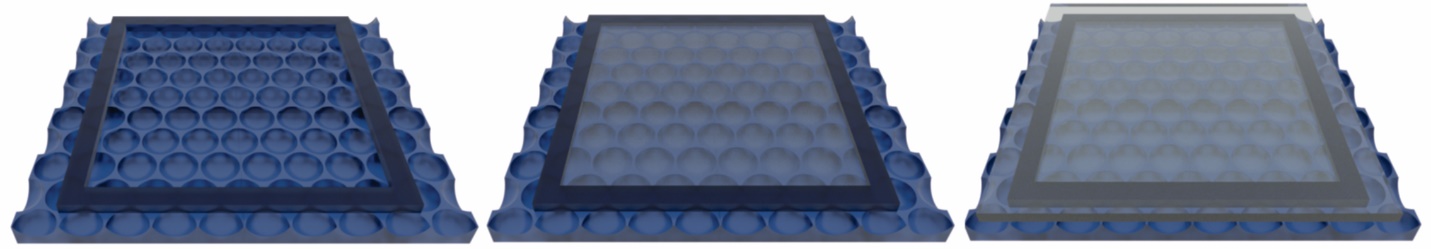

S.F. 2. Methodology for Raman measurements of bioparticles Firstly, 2 µl of sample were placed inside the PDMS chamber, placed on a nano-bowl SERS chip, and then sealed with a glass coverslip. The sealed solution was further placed under the Raman microscope for spectrum acquisition.

**Supplementary Section: 4**

**Bright field video on trapping of liposome:**

The EPE liposome of 143 nm mean diameter was captured inside the nano-bowl cured from 1 µm PS beads stamp. The methodology was followed the same as described in the supplementary section 3.

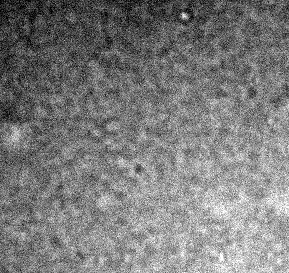

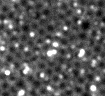

Left: S.M. 1 Lipoid free floating on flat PDMS, Right: S.M. 2 Lipoid trapping using structured PDMS video link

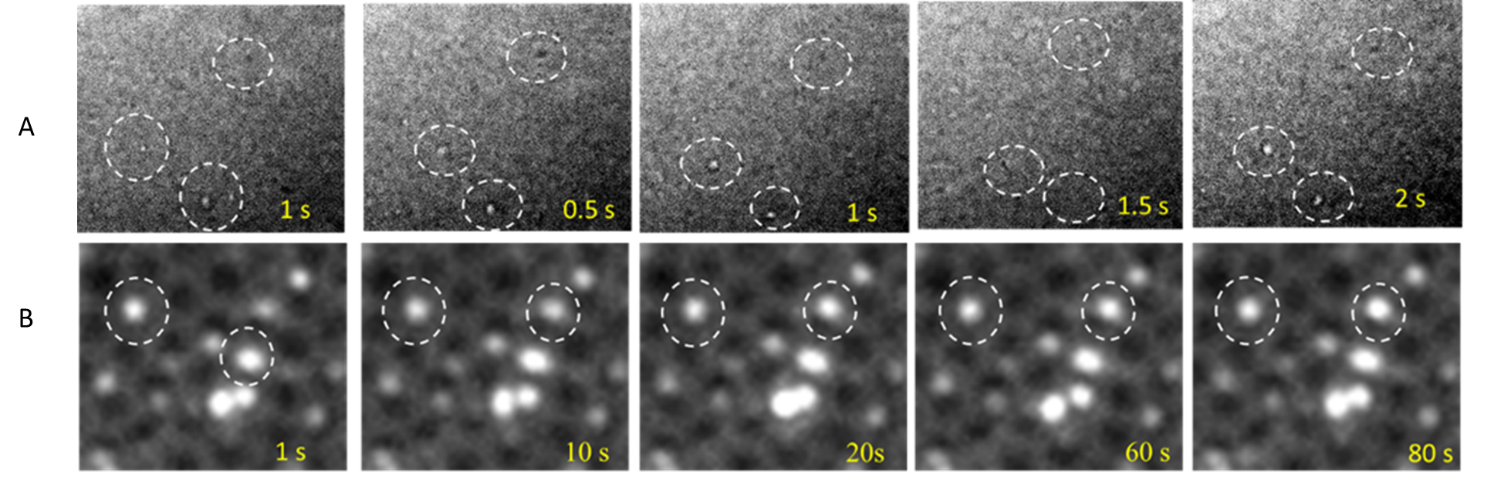

´S.F. 3 (a) Liposomes on row A (flat PDMS) and row B (nano-bowl) at various time intervals. Row B shows the lipoid trapped in the nano-bowl for longer duration, while row B shows the liposomes floating on the flat PDMS can move randomly due to their Brownian motion. The associated movie file S.M. 1 and S.M.2.

**Supplementary Section: 5**

**5.1 Raman and SERS spectra of lipids**

Raman spectra of SPC, EPE-liposomes, HAP1 and HAP1 F3KO derived EVs were collected considering a 2µL drop of solution inside a PDMS chamber, placed on flat Si wafer. The solution was further sealed using glass coverslip from top. The spectrum was acquired at a power of 60 mW, and integration time of 60 sec for 2 acquisitions.

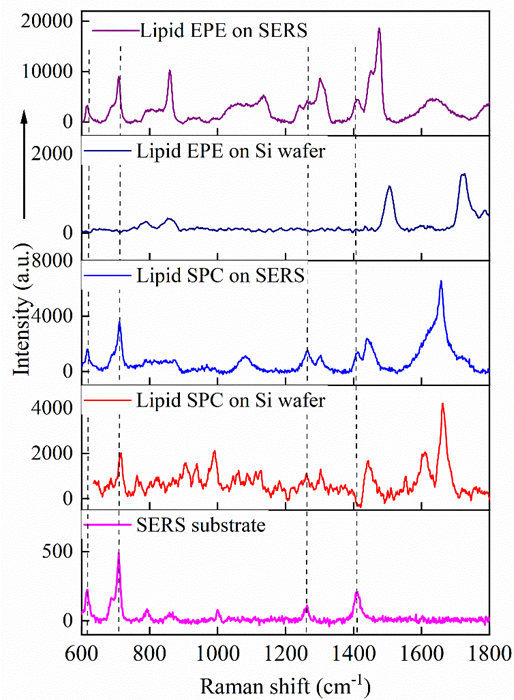

S.F. 4 Normal Raman and SERS spectra of liposomes. The PDMS based SERS substrate exhibits two prominent peaks at 709 cm-1 and 1411 cm-1 respectively. The normal Raman spectra of lipoid SPC exhibits peak at 1610 cm-1 and 1661 cm-1, respectively. Other signatures remain noisy. The normal Raman spectrum of EPE lipid exhibits peaks at 1345 cm^-1^, 1455 cm^-1^, and 1581cm^-1^ respectively. Both lipids exhibit enhanced and distinct peaks on SERS substrates. The Raman spectra were collected using 60 mW power, 60 sec integration time with 2 acquisitions, while the SERS spectra were acquired using a 10 mW laser power, 5 sec integration time with single acquisition. The dotted lines represent peak due to PDMS background.

**5.2 Raman and SERS spectra of EVs**

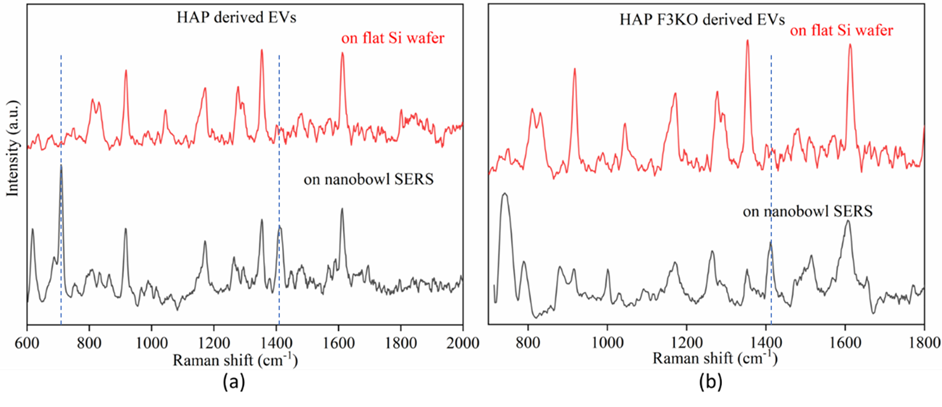

S.F. 5 Normal Raman and SERS spectrum of EVs isolated from HAP and HAPF3KO cells. The Raman and SERS spectra matches well for both EVs. The SERS enhancement of EVs was found to be less than the enhancement of liposomes samples. The Raman spectra were collected using 60 mW power, 60 sec integration time with 2 acquisitions, while the SERS spectra were acquired using a 30 mW laser power, 5 sec integration time with single acquisition. The dotted lines represent peak due to PDMS background.

**Supplementary Section :6**

**Reproducibility of SERS spectra**

The SERS spectrum reproducibility was checked for several independent measurements. The peaks are occurring at the same wavenumber value, and the intensity variation is calculated to be less than 10%.

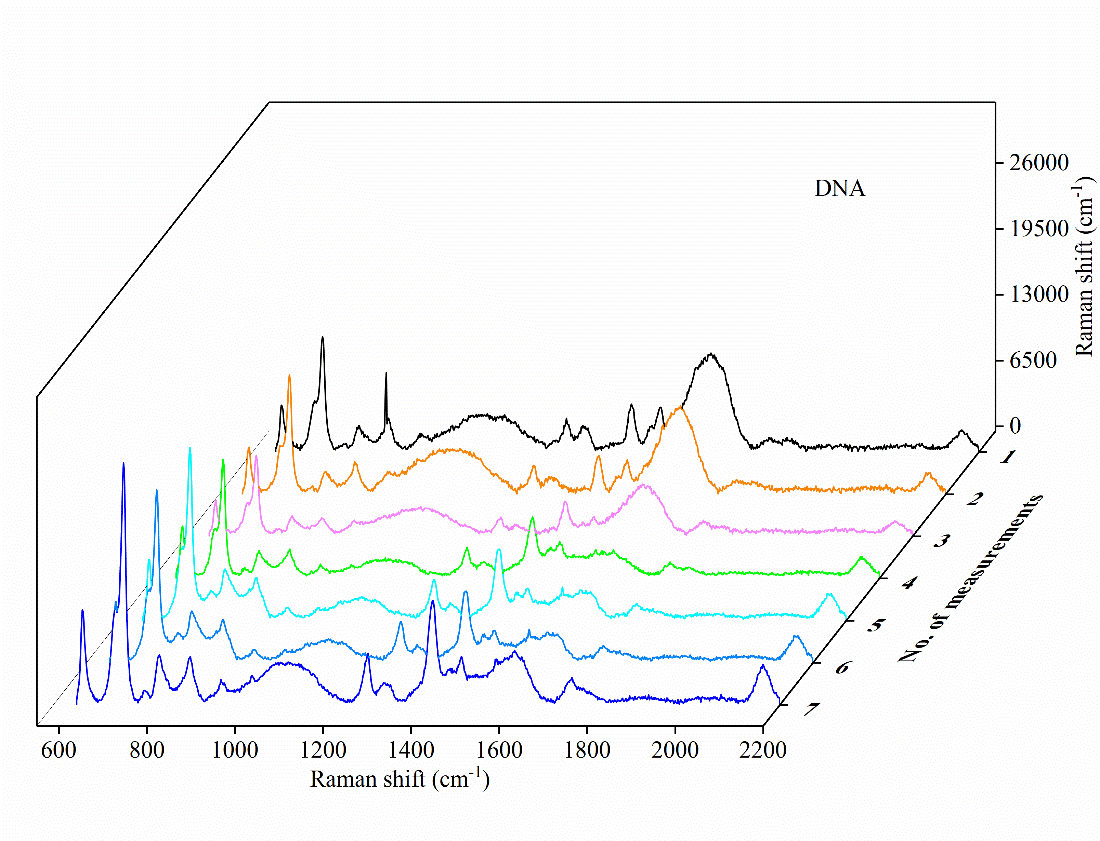

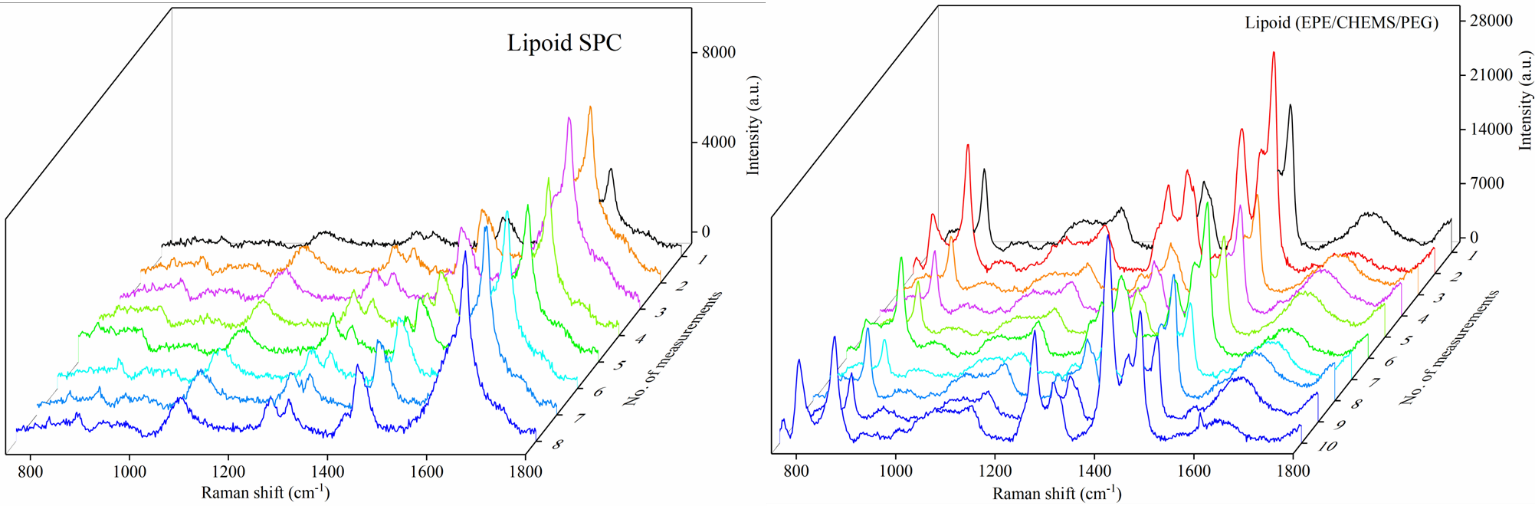

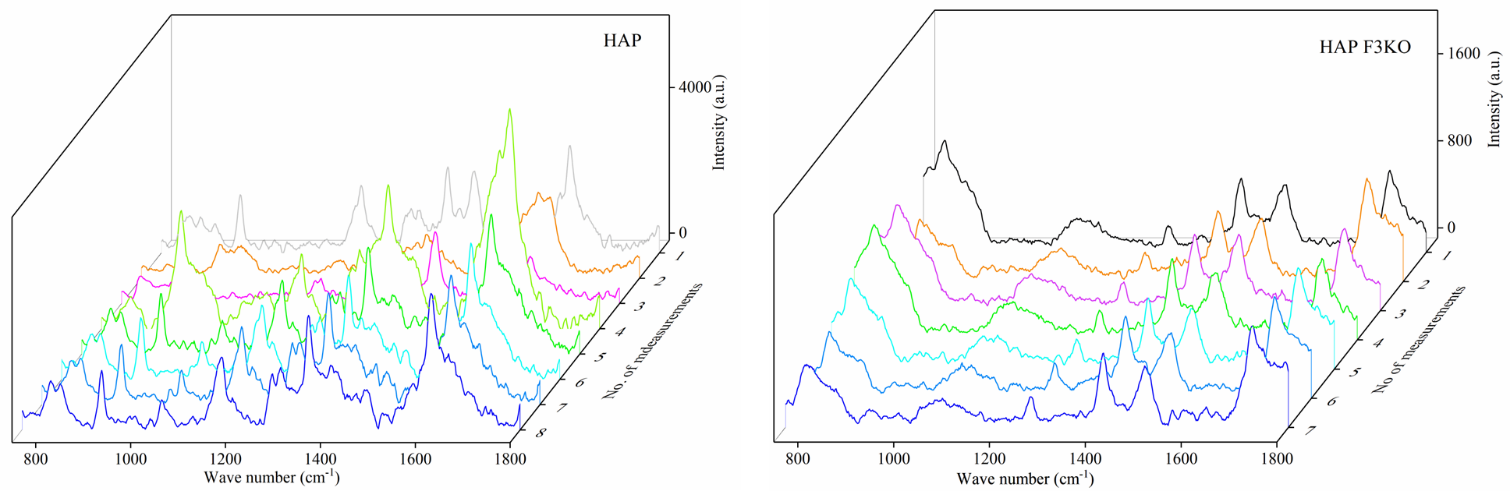

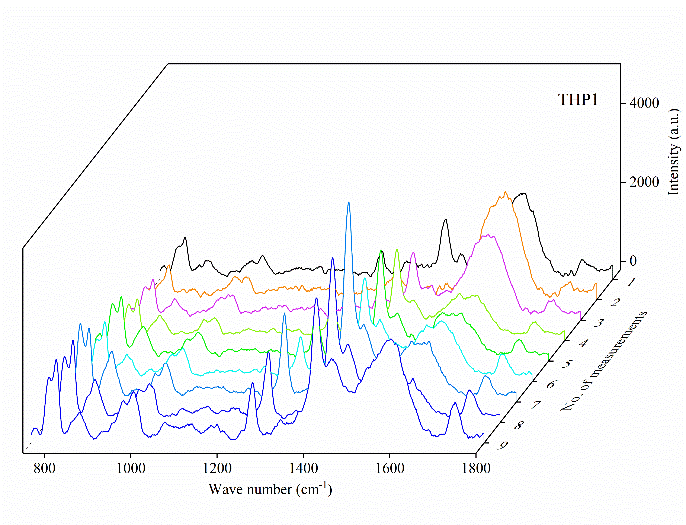

S.F. 6 SERS spectrum reproducibility for DNA, Liposomes, and EVs.
